# Single-cell trajectory inference guided enhancement of thyroid maturation *in vitro* using TGF-beta inhibition

**DOI:** 10.1101/2021.01.18.427103

**Authors:** Mírian Romitti, Sema Elif Eski, Barbara Faria Fonseca, Sumeet Pal Singh, Sabine Costagliola

**Author notes:** Equal Contribution (Mírian Romitti, Sema Elif Eski).

## Abstract

The thyroid gland regulates metabolism and growth via secretion of thyroid hormones by thyroid follicular cells (TFCs). Loss of TFCs, by cellular dysfunction, autoimmune destruction or surgical resection, underlies hypothyroidism. Recovery of thyroid hormone levels by transplantation of mature TFCs derived from stem cells *in vitro* holds great therapeutic promise. However, the utilization of *in vitro* derived tissue for regenerative medicine is restricted by the efficiency of differentiation protocols to generate mature organoids. Here, to improve the differentiation efficiency for thyroid organoids, we utilized single-cell RNA-Seq to chart the molecular steps undertaken by individual cells during the *in vitro* transformation of mouse embryonic stem cells to TFCs. Our single-cell atlas of mouse organoid systematically and comprehensively identifies, for the first time, the cell types generated during production of thyroid organoids. Using pseudotime analysis, we identify TGF-beta and planar-cell polarity (PCP) pathways as regulators of thyroid maturation *in vitro*. Using pharmacological manipulation of TGF-beta pathway, we improve the level of thyroid maturation, in particular the induction of *Nis* expression. This in turn, leads to an enhancement of iodide organification *in vitro*, suggesting functional improvement of the thyroid organoid. Our study highlights the potential of single-cell molecular characterization in understanding and improving thyroid maturation and paves the way for identification of therapeutic targets against thyroid disorders.

## 1 Introduction

### 1.1 Efficient generation of functional organs *in vitro* is pre-requisite for cell replacement therapy

Replacement of damaged or dysfunctional organs by *in vitro* generated tissues is a promising avenue for regenerative medicine. Cell replacement therapy has provided encouraging results against multiple degenerative diseases, including retinal degeneration (1), diabetes (2) and Parkinson’s disease (3). The progress has been enabled by the tremendous advances in development and directed differentiation of pluripotent stem cells -- embryonic stem cells (ESCs) and induced pluripotent stem cells (iPSCs). Notably, engineering of pluripotent stem cells to lack components of HLA (human leukocyte antigen), factors that are recognized by the immune system, provides a universal donor material for potential treatment of auto-immune disorders, such as Type 1 diabetes (4,5). The vast potential of cell replacement therapy is, however, hindered by the ability to mimic organ physiology *in vitro*. Production of patterned organs containing mature, functional cells at high efficiency and purity remains a major bottleneck.

*In* vitro generation of organs with natural architecture and physiology, called organoids, depends on recapitulation of regulatory steps involved in cell fate decision. Though the study of embryonic development has enriched our understanding of the differentiation process, our knowledge of gene dynamics during lineage specification and maturation remains incomplete. Recently, the utilization of single-cell RNA-Sequencing (scRNA-Seq) has allowed unbiased evaluation of the differentiation process (6-8). scRNA-Seq, by profiling the gene expression levels in individual cells, enables segregation of a heterogeneous cell-population undergoing asynchronous differentiation, thereby allowing identification of signaling pathways active at specific stages of differentiation. Modulation of the identified signaling pathways, particularly those that correlate with terminal differentiation and maturation, can improve the *in vitro* protocol for generation of functional tissues.

Here, we combine scRNA-Seq with an organoid model of the thyroid gland to discern the signaling pathways regulating thyroid differentiation and utilize the information to improve the protocol by modulation of an identified pathway. For this, we exploit a method our group had previously developed for *in vitro* generation of functional thyroid (9). Our thyroid organoid protocol generates 3D, patterned tissue from mouse ES cells. The differentiated organ was shown to be capable of rescuing a mouse model of complete thyroid loss, underscoring the functional activity of the *in vitro* generated organ. Though variations of the protocols have been since published (10), two significant shortcomings exist in the field: 1. we lack a comprehensive understanding of the distinct cell lineages present in the organoids, particularly non-thyroid cell types; and 2. we lack systematic analysis of gene regulatory networks underlying the differentiation process. We address these gaps by analysis of mouse thyroid organoid differentiation at single-cell resolution. We comprehensively identify the cell types present in the organoid, decipher the pathways regulating thyroid maturation and finally, utilize this information to improve the efficiency of *in vitro* differentiation protocol.

### 1.2 Brief introduction of thyroid function and *in vivo* development

Thyroid gland is an endocrine organ responsible for synthesis, storage and secretion of thyroid hormones (triodothyronine (T3) and thyroxine (T4)). The functional unit of the gland are hollow, spherical follicles that store immature thyroid hormone in a colloidal form within the follicle lumen. The follicles are composed of polarized epithelial thyroid follicular cells (TFCs), also known as thyrocytes. Thyrocytes express Thyroglobulin (*Tg*), which serves as the precursor to thyroid hormone. Production of thyroid hormones from *Tg* requires a complex series of reactions that involves bidirectional transport to and from the lumen (recently reviewed in (11,12)). This process is orchestrated by contributions from multiple genes, including thyroperoxidase (*Tpo*) and sodium/iodide symporter (*Nis* / *Slc5a5*).

*In vivo* the gland develops from the anterior foregut endoderm. The process involves recruitment of a group of endodermal cells to thyroid fate, in an event called “specification” or “determination”, which at the molecular level, is characterized by the co-expression of *Nkx2-1* (TTF-1), *Foxe1* (TTF-2), *Pax8* and *Hhex* transcription factors (13-18). Subsequently, thyroid progenitors undergo a series of events which comprise cell proliferation, invasion of the surrounding mesenchyme, migration, thyroid lobes enlargement and consequent appearing of follicular organization (19,20). The thyroid differentiation program is complete when the gland reaches its final location and TFCs express a series of specific proteins which are essential for thyroid hormonogenesis. The thyroid markers present a temporal pattern of expression, in which thyroglobulin (*Tg*), thyroperoxidase (*Tpo*), and TSH receptor (*Tshr*) are detected initially (17), followed by sodium/iodide symporter (*Nis*) (21) and *Duox1/2* expression (22), resulting in thyroid hormone synthesis, storage and secretion.

*In vitro* differentiation of thyroid gland from pluripotent stem cells is capable of generating polarized, spherical follicles that synthesize thyroid hormone (9). However, it remains unknown if the sequential expression of genes observed *in vivo* is recapitulated *in vitro*. Here, using single-cell RNA-Seq of mouse thyroid organoid model, we reconstruct the gene expression dynamics during *in vitro* differentiation and show that it matches well with the expectation from studies on *in vivo* differentiation of the thyroid gland.

## 2 Results

### 2.1 Single-cell RNA-Seq of mouse organoid

Our previously published protocol was used to generate mouse thyroid organoid from mouse embryonic stem cells (mESCs) (9). Here, we utilized mESCs that had been engineered to express three constructs (Figure 1A): 1. Rosa26 locus driven Tet-On transcription factor (rtTA: reverse tetracycline-controlled transactivator), 2. At the Hypoxanthine phosphoribosyltransferase (HPRT) locus was inserted Tet-Responsive Element (TRE) driven bicistronic construct containing Nkx2.1 and Pax8, and 3. *Tg* promoter driven EGFP. The three constructs allow doxycycline-induced expression of exogenous *Nkx2*.*1* and *Pax8* (referred as exNkx2.1_Pax8 henceforth) and monitoring of the thyroid lineage by Tg-driven EGFP expression.

**10.1 Figure 1:**
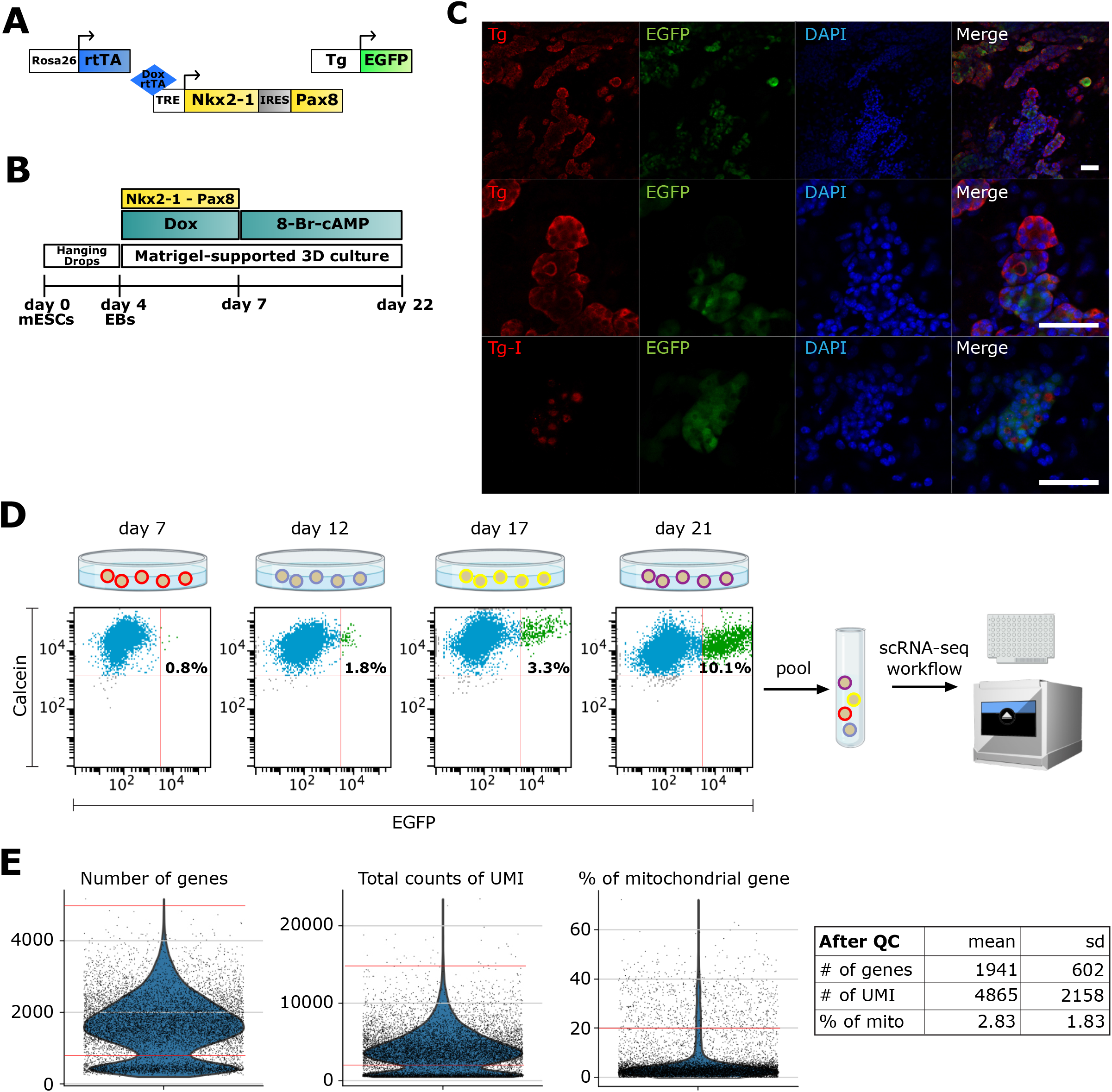
Single-cell RNA-Seq of mouse organoid. **(A)** Schematic of the three constructs used in the protocol: Rosa26 locus driven reverse tetracycline-controlled transactivator (rtTA), Tet-Responsive Element (TRE) driven *Nkx2*.*1* and *Pax8*, and *Tg* promoter driven EGFP. **(B)** Schematic of differentiation protocol for mESC-derived *in vitro* thyroid organoid. **(C)** Immunofluorescence images displaying differentiated organoids marked with Thyroglobulin (*Tg*) (red), EGFP (green) and DAPI (blue). Middle and bottom rows are zoomed in images of upper rows (Scale bars: 50 µm). **(D)** Cells from differentiated organoids at days 7, 12, 17 and 21 were labeled with calcein (live cell marker) and FACS-sorted for scRNA-seq analysis. Percentage of EGFP+ thyrocytes are noted. For downstream analysis, EGFP+ and EGFP-cells (1:1 ratio) from day 12, 17 and 21 were pooled into a single tube and sequenced using 10X Genomics platform. **(E)** Violin plots showing the quality control parameters for cells profiled using scRNA-seq. Number of genes, total counts of UMI and the percentage of mitochondrial genes were utilized for quality control. Red lines on violin plots depict the cutoffs for filtered cells. After filtration, cells from scRNA-seq had a mean ± standard deviation of 1941 ± 602 genes, 4865 ± 2158 UMIs and 2.83 ± 1.83 % of mitochondrial gene content.

Thyroid organoids are generated from mESCs in a multi-stage protocol (9) (Figure 1B). Briefly, the mESCs are differentiated into embryoid bodies (first step towards differentiation, not endoderm/thyroid specific) by day 4, using the hanging drop technique, upon which the cells are embedded in matrigel and treated with doxycycline for three days to induce the expression of exNkx2.1_Pax8 until day 7. At this stage, cells are treated with thyrotropin hormone (or cAMP) until day 22 or longer. Our previous work had established that by day 22, thyroid follicular cells are fully mature, expressing thyroid transcription factors, *Nkx2*.*1, Pax8, Hhex* and *Foxe1*, as well as functional markers such as *Tg* (Figure 1C), *Tshr, Tpo* and *Nis*. Moreover, thyroid cells self-organize into three-dimensional follicular structures containing iodinated thyroglobulin (Tg-I) in the lumen (Figure 1C).

To obtain a cellular resolution understanding of the multi-stage process, we performed single-cell RNA-Seq (scRNA) from cells collected at four stages of the protocol (Figure 1D): day 7 (end of doxycycline-based induction of exogenous *Nkx2*.*1* and *Pax8*), day 12 (early development), day 17 (advanced differentiation), and finally day 22 (functional thyroid follicles). To avoid batch effects, we combined 4,000 cells from each stage, 50% of them were GFP+ and 50% GFP- and profiled them using droplet based scRNA-Seq from 10x Genomics. The droplet-based scRNA-Seq encapsulates each cell into a microfluidic based nanoliter-sized reaction chamber, where the cells are lysed, and mRNA reverse transcribed to cDNA for next-generation sequencing. Notably, during reverse transcription, each mRNA molecule is tagged with a unique barcode, referred to as Unique Molecular Identifier (UMI) (23), which reduces the impact of cDNA amplification bias. Upon sequencing the library to a depth of 105 million reads and demultiplexing, we obtained profiles from 9,904 cells. The profiled cells underwent quality control on the basis of UMI, number of genes and percentage of mitochondrial reads (Figure 1E), yielding a total of 7,381 filtered cells for downstream analysis.

### 2.2 Identification of cell types present in thyroid organoid

To visualize distinct cell types in mESC-derived thyroid organoids, we projected the scRNA-seq data onto two dimensions using a non-linear dimensionality reduction technique, UMAP (24), and performed unsupervised graph-based cell clustering (Figure 2A). For each cluster, marker genes were identified using Scanpy (25) (Supplementary Table 1). Marker genes display statistically significant enrichment in expression levels within a particular cluster. Using the list of marker genes, we annotated cell types with literature survey. Overall, we identified nine clusters for mESC-derived thyroid organoid model (Figure 2A). Of interest, we could identify a cluster of thyroid follicular cells containing 2,914 cells that display expression of genes involved in thyroid gland development and thyroid hormone production, including *Tg, Pax8, Foxe1, Hhex* as well as EGFP driven from Tg promoter (Figure 2B, C).

**10.2 Figure 2:**
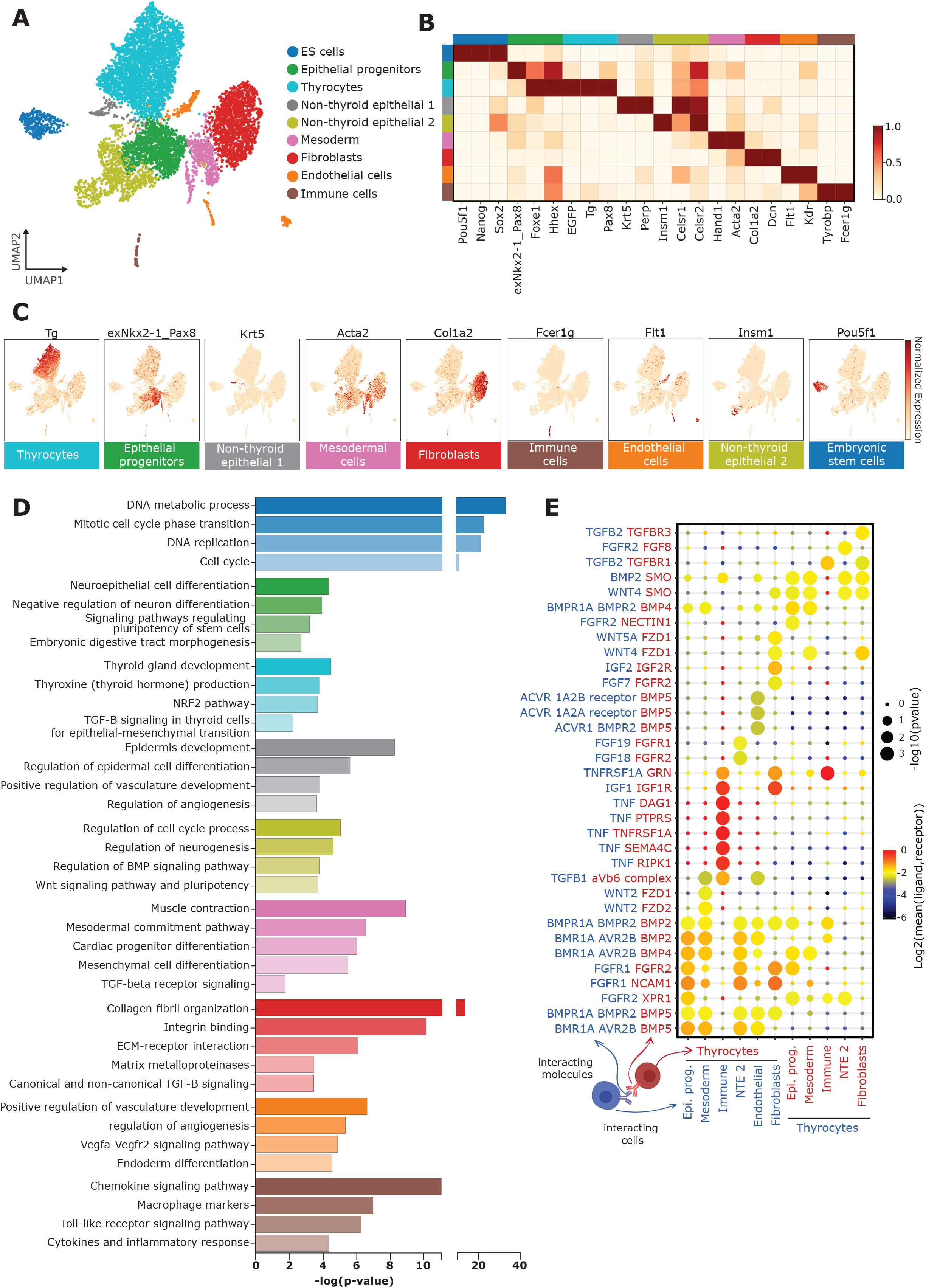
Characterization of cell types present in thyroid organoids. **(A)** Unsupervised clustering of *in vitro* thyroid organoid model. Each cluster is represented by a specific color. **(B)** Gene-expression matrix plot for selected marker genes for each cell cluster. Rows depict cell clusters, while columns depict genes. The intensity of color in each square indicates mean expression within the cluster. **(C)** UMAP expression plots of representative marker genes for thyrocytes (*Tg*), epithelial progenitors (exNkx2-1_Pax8), non-thyroid epithelial cells 1 (*Krt5*), mesodermal cells (*Acta2*), fibroblasts (*Col1a2*), immune cells (*Fcer1g*), endothelial cells (*Flt1*), non-thyroid epithelial cells 2 (*Insm1*) and embryonic stem cells (*Pou5f1*). Color intensity indicates normalized expression of respective genes. **(D)** GO term analysis for differentially expressed genes. Color of the bar plots correspond to the color of respective cell cluster. **(E)** CellPhoneDB-generated dot plot showing potential ligand-receptor pairs between thyrocytes and epithelial progenitors, mesodermal cells, immune cells, non-thyroid epithelial cells 2 (NTE 2), endothelial cells, fibroblasts. Each dot’s color intensity correlates with log2 of mean expression of ligand-receptor pair between the two clusters, while size of the dots indicates −log of p-value.

As described in our previous study where we established the mouse thyroid organoid (9), the organoid model generates up to 60% of the cells that express endogenous Nkx2.1 and Pax8. The identity and molecular nature of the remaining cells has so far remained undocumented. Using literature survey of marker genes, we annotated the non-thyroid cells present in the organoid model (Figure 2A - C). Based on cluster-specific signature genes, we identified 282 embryonic stem cells expressing stemness markers such as *Pou5f1 (Oct4), Nanog, Sox2*; 209 endothelial cells enriched in markers such as *Flt1, Kdr, Cd34, Sox17*; 42 immune cells with known markers such as *Tyrobp, Fcer1g, Csf1r, Lyz*; 502 mesodermal cells expressing mesoderm lineage marker genes such as *Hand1, Hand2, Gata6, Acta2*; 1363 fibroblast cells with expression of pan-fibroblast markers such as *Col1a2, Col1a1, Dcn*; 76 non-thyroid epithelial cells 1 with epithelial cell specific marker gene expression such as *Krt5, Krt15*; 910 non-thyroid epithelial cells 2 with expression of markers such as *Insm1, Celsr1, Celsr2*; and finally 1083 epithelial progenitor cells which express a mixture of early thyroid markers and epithelial specific markers in addition to exogenous bicistronic *Nkx2-1/Pax8* construct (exNkx2-1_Pax8) expression.

To gain further insights into the gene expression profiles of different clusters, we performed GO Term analysis for marker genes (Figure 2D). This revealed involvement of distinct metabolic and biological processes to each cell type. Of note, thyrocytes showed an enrichment of genes involved in endocrine system development, thyroid gland development, thyroxine (thyroid hormone) production, NRF2 pathway, TGF-beta signaling pathway and Wnt signaling pathway. Epithelial progenitors were enriched for genes involved in regulation of cell cycle process, regulation of neurogenesis, BMP and Wnt signaling pathways and pluripotency. Fibroblast cluster displayed enrichment of genes involved in ECM-receptor expression, collagen fibril organization, matrix metalloproteinases and canonical and non-canonical TGF-beta and Wnt signaling, whereas mesoderm cluster showed enrichment of genes involved in vasculature development, endoderm differentiation and VEGFR signaling pathway.

### 2.3 *In silico* connectome for thyroid organoid model

Our single-cell atlas provides a molecular map for the cells present in the thyroid organoid. We exploited this map to develop a connectome showcasing potential paracrine signaling active in the organoid model. For this, we utilized CellPhoneDB (26) which predicts interaction between two cell types using a database of ligands and their corresponding receptors. Additionally, as we had mixed multiple stages of organoids into one sequencing reaction, remaining non-induced pluripotent stem cells were present in the dataset (Fig. 2A). To explore possible molecular interactions which could be relevant in *in vivo* context, we excluded from connectome analysis the ESC cluster and considered molecular interaction between the remaining cell types. Thus, we considered interaction between thyrocytes, epithelial progenitors, endothelium, mesoderm, fibroblasts, non-thyroid epithelial 2 and immune clusters. With this, we could detect a total of 136 significant interactions (Figure 2E) (Supplementary Table 2). These interactions were enriched for TGF-beta, Wnt, BMP and FGF signaling pathways.

Several interactions identified with the connectome caught our attention, particularly considering the role of the aforementioned signaling pathways in thyroid specification (27). For instance, interaction between epithelial progenitors and thyrocytes were predominantly enriched for BMP signaling pathway. Explicitly, *Bmp2* and *Bmp4* ligands expressed by thyrocytes bind to *Bmpr1a* and *Avr2b*, respectively, which are expressed by epithelial progenitors. Similar interactions were also observed between thyroid and non-thyroid epithelial 2 or endothelial cluster. Ligand-receptor pairs showed a possible interaction between immune cells and thyrocytes, with immune cells expressing *Tnf* and thyrocytes expressing *Sema4c, Tnfrsf1a*. Further, the interaction between mesodermal cells and thyrocytes is enriched in BMP and Wnt pathways. Specifically, *Bmp5* from thyrocytes finds an interacting partner in *Bmpr1a* and *Bmpr2* in mesoderm, while *Wnt2* expressed by mesodermal cells could potentially signal via *Fzd1* and *Fzd2* expressed by thyrocytes. Finally, the TGF-beta ligand *Tgfb2* is expressed in thyrocytes signals through the TGF-beta receptors *Tgfbr1* and *Tgfbr3* expressed by fibroblasts.

### 2.4 Pseudotime analysis reveals gene expression dynamics during thyroid differentiation

Next, we focused on the thyrocyte cluster to gain detailed information on the factors regulating thyrocyte morphogenesis. For this, we noted that while thyrocytes uniformly expressed *Thyroglobulin* (*Tg*), they did not uniformly express genes related to thyroid functionality (Figure 3A-C). A proportion of thyrocytes expressed *Tpo* and *Nis* / *Slc5a5*. We labeled *Tg*+ *Tpo*-*Slc5a5*-cells as immature thyrocytes and *Tg*+ *Tpo*+ or *Slc5a5*+ cells as mature thyrocytes. Mature thyrocytes constituted 7% of all thyrocyte population. It is worth noting that immature thyrocytes express *Pax8, Foxe1* and *Hhex* (Figure 3B, C), and while the expression of *Tshr* was low, it was more uniform than the expression of *Tpo* or *Slc5a5* within the thyrocyte cluster (Figure 3C).

**10.3 Figure 3:**
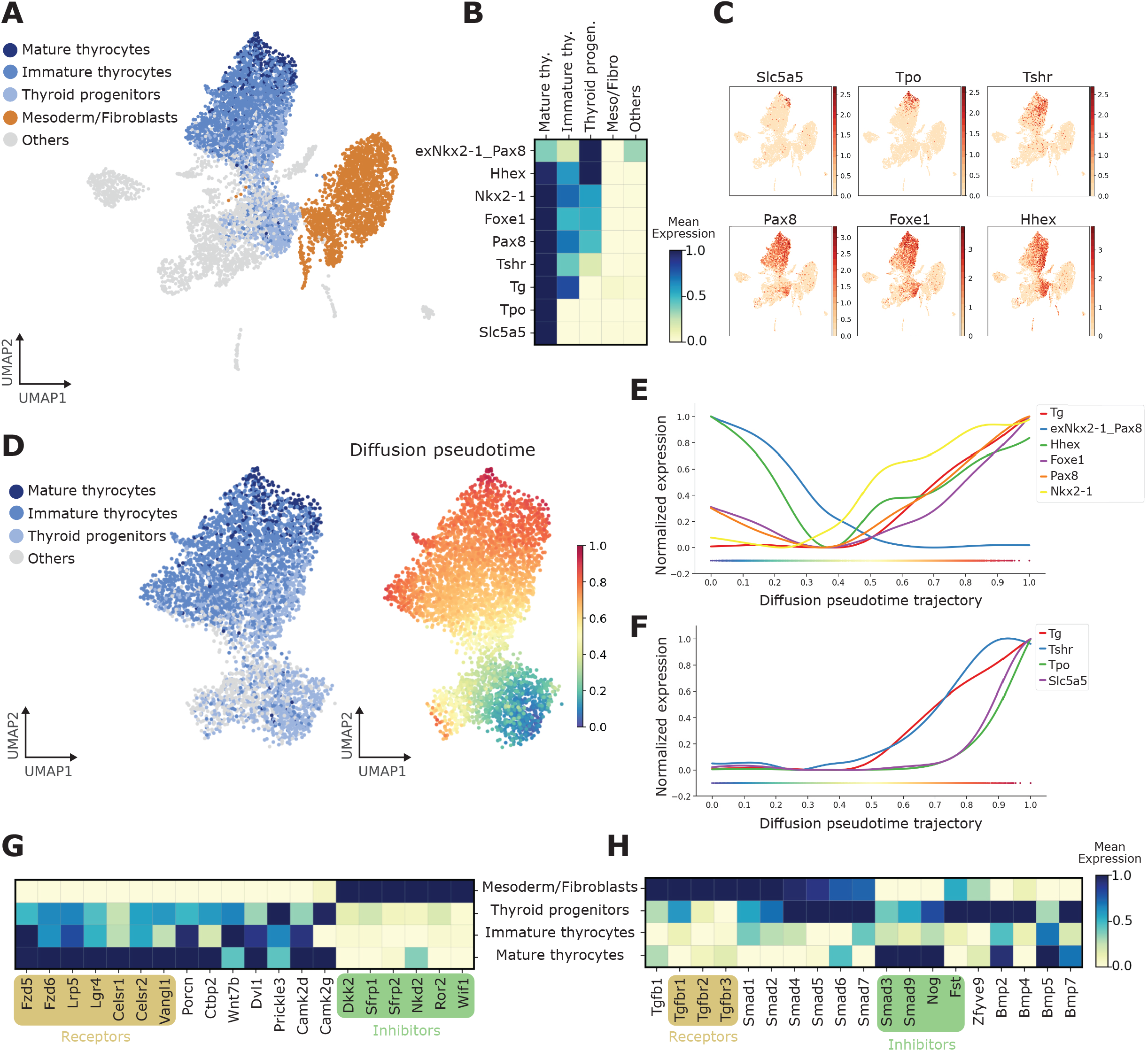
Identification of pathways dynamically regulated during TFCs maturation. **(A)** UMAP displaying subclusters within the thyroid lineage (thyroid progenitors, immature and mature thyrocytes). Remaining cells are labeled as ‘other’. **(B)** Gene-expression matrix plot for thyroid markers: *Hhex, Nkx2-1, Foxe1, Pax8, Tshr, Tg, Tpo, Nis* / *Slc5a5* and exogenous Nkx2-1_Pax8 in subclusters of thyroid lineage and other cell groups. Color intensity in each square represents mean expression in the corresponding cluster. **(C)** UMAP overlaid with gene expression plots for thyrocyte markers. Color indicates normalized expression. **(D)** Diffusion pseudotime analysis of thyrocyte lineage. (Left) UMAP displaying the subclusters within the thyroid lineage; (Right) UMAP overlaid with pseudotime. Color in pseudotime plot indicates the ordering of the cells. **(E-F)** Expression trends of specific thyroid markers along pseudotime trajectory **(G - H)** Gene-expression matrix plot for selected genes involved in Wnt/PCP pathway (G) and TGF-beta pathway (H). The gene list includes receptors and inhibitors. Color intensity in each square indicates mean expression in the corresponding subpopulation.

Further, to extend the analysis to the cellular source of thyrocytes, we included cells labeled as ‘Thyroid Progenitors’, as defined by cells that co-express exogenous or endogenous *Nkx2*.*1* and *Pax8*, along with *Hhex* and *Foxe1*, but lack *Tg* expression (Figure 3B). This annotation was based on the expression of known marker genes for thyroid progenitors (13). Remaining cells in the atlas were labeled as ‘Others’.

By combining thyroid progenitors with thyrocytes segregated by maturation status, we could obtain a cellular progression recapitulating thyroid differentiation and maturation.

Using the progressive nature of thyrocyte morphogenesis, we constructed an *in silico* temporal ordering of the thyrocyte cluster. In this, a trajectory of cellular progression from one state to another is predicted based on similarity in gene expression (28). Using pseudotime analysis, we estimated the transition from thyroid progenitor to mature thyrocytes (Figure 3D) and utilized this to generate the trend of gene expression along the trajectory (Figure 3E-F). With this, we observed that a decline in exogenous *Nkx2-1*/*Pax8* expression coincided with an increase in endogenous *Nkx2-1* and *Pax8* expression, suggesting the presence of a positive feedback loop, as has been demonstrated previously (9,29,30). Interestingly, the trend of *Hhex* expression closely follows the dynamics of *Nkx2-1* and *Pax8*, irrespective of exogenous or endogenous source of transcription factors (Figure 3E). Moreover, expression of *Foxe1* correlated with the dynamics in *Tg* expression (Figure 3E). With regard to genes related to thyroid maturation, *Tg* and *Tshr* display similar trends, while the expression of *Slc5a5* and *Tpo* increase steeply at the end of pseudotime trajectory (Figure 3F).

It is worth pointing out that the pseudotime analysis indicates two possible differentiation branches originating from progenitors (Figure 3D). One of the branches moves towards non-thyroid lineages, suggesting that part of the cells expressing doxycycline induced exogenous *Nkx2-1*/*Pax8* can differentiate to non-thyrocyte lineages. This potentially explains the presence of non-thyroid epithelium in the thyroid organoid model and suggests presence of additional factors that specifically restricts the progenitors to thyroid lineage.

### 2.5 TGF-beta and planar cell polarity display dynamic regulation during thyroid maturation

To identify key signaling pathways regulating the maturation of thyrocyte, we utilized trajectory analysis to identify genes and pathways that correlate with the progression in thyroid lineage. Further, we compared the gene expression with mesodermal/fibroblast lineage, which acted as a negative control (Fig. 3G, H). With this, we identified two pathways being pivotal in thyrocyte lineage progression: Non-canonical Wnt / Planar Cell Polarity (PCP) pathway (Figure 3G) and TGF-beta signaling pathway (Figure 3H). Specifically, positive regulators for PCP pathway, such *as Lrp5, Porcn, Fzd5, Dvl1, Celsr1* and *Vangl1*, are enriched in thyrocyte lineage and increased during maturation. In contrast, inhibitors of PCP pathway, such as *Dkk2, Sfrp1, Sfrp2, Wif1*, are exclusively expressed in mesoderm/fibroblasts. Further, fibroblasts express higher levels of TGF-beta receptors, *Tgfbr1, Tgfbr2, Tgfbr3*, as well as downstream activators, including *Smad1* and *Smad2*. Thyrocytes, on the other hand, are enriched for TGF-beta antagonists such as *Smad3, Smad9, Noggin and Fst*. Notably, expression of TGF-beta pathway inhibitors is prominently high in early thyrocytes, whilst decreasing through the maturation. Overall, these results suggest that dynamics in Wnt/PCP and TGF-beta pathways may play a decisive role in thyrocyte differentiation and maturation, possibly in interaction with mesoderm/fibroblasts.

### 2.6 Inhibition of TGF-beta pathway improves the efficiency of thyroid maturation *in vitro*

In order to access the scRNAseq-detected inhibitory effect of TGF-beta pathway on maturation of thyrocytes, we treated our organoids with TGF-beta receptor inhibitor, SB431542 (10µM), in addition to cAMP, for seven days (From day 22 to day 29; Figure 4A). Interestingly, we observed that TGF-beta inhibition results in significant induction of *Tg* and *Pax8* gene expression (p<0.05; Figure 4B). Moreover, TGF-beta inhibition seems to have a pronounced effect on *Nis* regulation, since SB431542 treatment resulted in more than 2-fold induction in the *Nis* expression level compared to the control (p < 0.0001; Figure 4B). *Nis* / *Slc5a5* transports iodide into thyrocytes. Thus, we hypothesized that an increase in *Nis* expression due to TGF-beta inhibition could improve the capacity of the thyrocytes to sequester iodide from the culture media. To test such improvement in thyroid function, we performed iodide organification assay. In this functional assay, we measured the uptake and incorporation of radioiodine into precipitable proteins. For this, organoids were incubated for 2 hours with ^125^I-supplemented medium prior to analysis. In line with our hypothesis, the inhibition of TGF-beta pathway increased the capacity of thyrocytes to uptake iodide (p < 0.001; Figure 4C), while also improving the efficiency of iodide-binding to Tg (p < 0.05; Figure 4D).Consequently, this leads to a higher percentage of cells promoting iodide organification (Figure 4E). In addition, immunofluorescence staining confirmed that the TGF-beta inhibition did not affect thyroid follicular organization and demonstrated specific *Nis* expression with Tg-I presence in the lumen space (Figure 4F).

**10.4 Figure 4:**
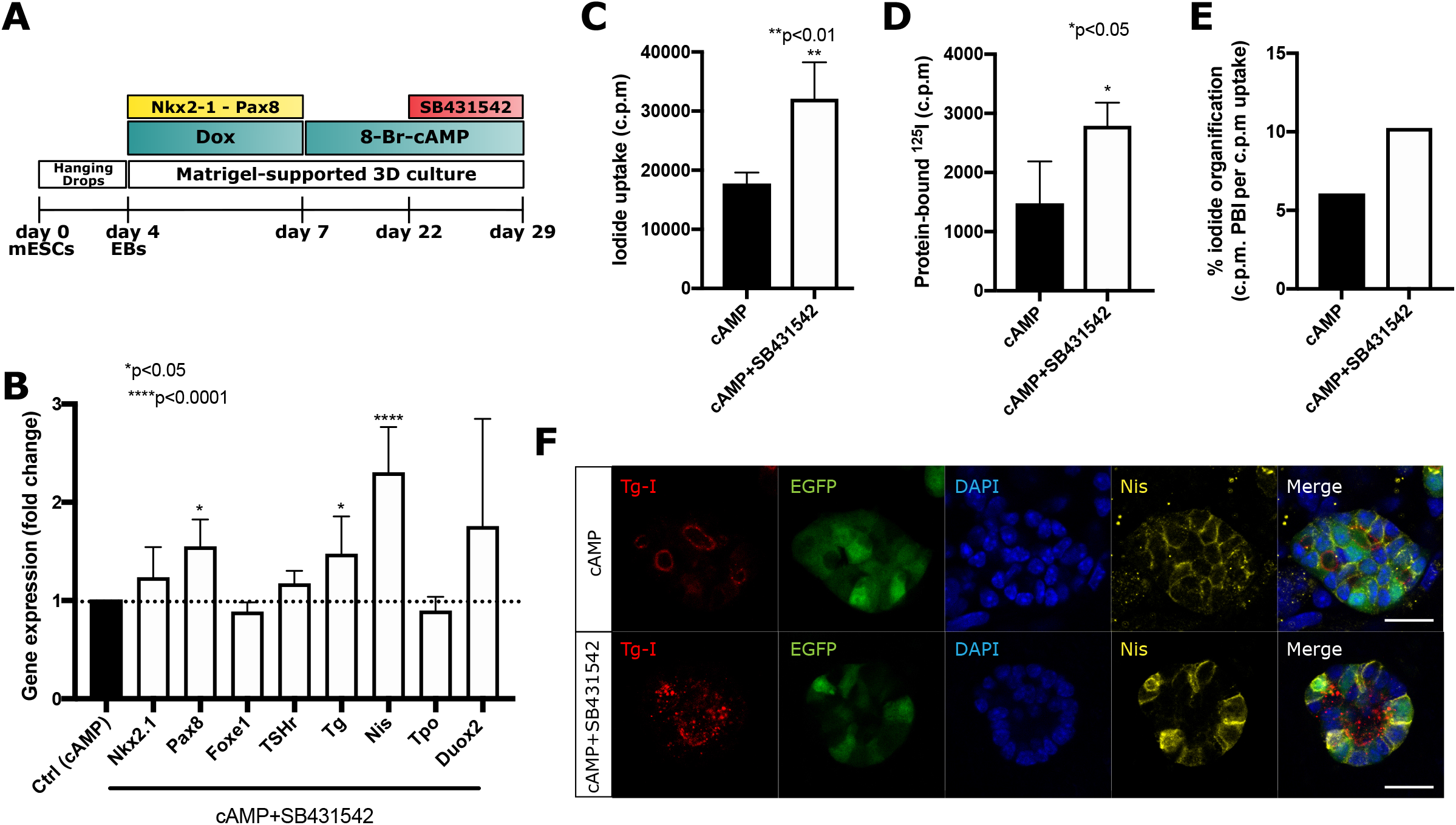
Enhancement of maturation efficiency by pharmacological inhibition of TGF-beta. **(A)** Schematic representing the thyroid differentiation protocol to test the impact of TGF-beta inhibition on thyroid organoid maturation. TGF-beta inhibition was induced by co-treatment with cAMP+SB431542 from day 22 to day 29. **(B)** qPCR analysis demonstrates significant upregulation in *Pax8, Tg* and *Nis* gene expression after 7 days of TGF-beta pathway inhibition. ^*^p<0.05,^****^p<0.001 by one-way ANOVA followed by Tukey’s test **(C - E)** Functional organification assay confirms significant improvement in 125I uptake (C) and protein-bound 125I (D) under SB431542 treatment, which results in a higher percentage of cells with capacity of 125I organification (E). ^*^*p< 0.01, ^*^p<0.05 by Student’s t-test. **(F)** Immunofluorescence staining demonstrating that SB431542 treatment does not affect thyroid follicular organization evidenced by *Nis* specific expression and Tg-I presence in the lumen space Scale bar: 20 µm.

## 3 Discussion

Substantial progress has been made in uncovering the molecular regulators of thyroid development and function (13,31); yet less than 5 % of congenital thyroid disorder cases have a known genetic cause (32,33). A better and wider characterization of the key regulators of thyroid differentiation is required to understand the development of thyroid disorders. Here, by taking advantage of unbiased single-cell RNA sequencing of our previously established mESC-derived thyroid organoid model, we have generated a molecular map of the thyroid gland lineage at single-cell resolution. Moreover, we identified some key signaling pathways driving thyroid cell differentiation. Specifically, our analysis revealed the involvement of TGF-B along with the non-canonical Wnt/PCP pathway in regulating thyroid gland lineage. Moreover, pharmacological manipulations of TGF-B pathway confirmed its inhibitory effect on thyroid maturation and function.

Thyroid-stimulating hormone (TSH) is the main primary physiological regulator of thyroid growth and function (34). Besides TSH/cAMP signaling, recent studies have suggested extrinsic regulators that tightly control folliculogenesis and maturation. Among them, multiple lines of evidence indicate a regulatory role of transforming growth factor-beta 1 (TGF-B1) and epidermal growth factor (EGF) ligands on thyroid cell proliferation, differentiation and function (35,36). Our observations on the inhibitory role of TGF-beta pathway on thyroid maturation fits well with the current literature, which underscores the capacity of our organoid model to mimic aspects of *in vivo* differentiation of the thyroid gland.

Further, mouse thyroid development occurs as a consecutive cascade of events which can be identified by the expression of specific thyroid markers (13). Taking this temporal gene expression into account, we could order individual thyrocytes along a linear differentiation trajectory (Figure 3D). The ordering allowed us to identify signaling pathways specifically enriched in progenitors versus mature thyrocytes (Figure 3E - H). For instance, BMP2 and BMP4 ligands, together with BMP-related Smad effectors, are enriched in thyroid progenitors in comparison with mature thyrocytes (Figure 3H). Accordingly, BMP pathway has been demonstrated to play a crucial role in thyroid specification and differentiation, *in vitro* and *in vivo* (10,27).

In addition to the study of TGF-beta and BMP pathway, we provide evidence demonstrating the enrichment of Wnt/PCP signaling components, both ligands and receptors, specifically in mature thyrocytes (Figure 3G). Notably, Wnt/PCP pathway plays a critical role in the differentiation and function of endocrine cells in the pancreas. Particularly, the pancreatic progenitors, which generate insulin-secreting pancreatic beta-cells, rely on PCP cues for establishing apico-basal polarity that is required for differentiation to endocrine lineage (37,38). Work from the group of Grapin-Botton has demonstrated the requirement for *Celsr2* and *Celsr3* in pancreas morphogenesis (37). Mice double mutant for *Celsr2* and *Celsr3* have a reduction in the number of beta-cells and are unable to maintain normal blood glucose levels. Further, they demonstrated that mislocalization of *Vangl2* from apical to basolateral membrane led to apoptosis of progenitor cells and a reduction in the size of pancreas. Notably, in our thyroid organoids, *Celsr1, Celsr2* and *Vangl1* expression increases during thyroid maturation (Figure 3G). Moreover, work from Lickert group demonstrated that the expression of *Flattop* (*Cfap126*), a PCP pathway effector gene, is able to segregate pancreatic beta-cells into proliferative and functional sub-populations (39). The role of Wnt/PCP in the endocrine pancreas, along with our data demonstrating its increase during *in vitro* maturation (Figure 3G), warrants a detailed analysis of Wnt/PCP pathway in the polarized thyroid follicular cells.

Besides the molecular characterization of the thyroid organoid, our study provides a map of the cell-types present in the culture. Notably, we observe presence of fibroblasts in the organoid model. Fibroblasts are part of the connective tissue present in the thyroid gland (40,41). Fibroblasts play an essential role in the physiology of an organ. For instance, fibroblasts have been shown to be critical for homeostasis of intestinal epithelial *in vivo* and in organoid cultures (42-44). Notably, sub-populations of intestinal fibroblasts express non-canonical Wnt ligands and BMP inhibitors (45), generating the necessary microenvironment for maintenance of stem cell niche. Further, remodeling of collagen by fibroblasts has been implicated in the progression of thyroid cancer (46). Thus, the presence of fibroblasts in the thyroid organoid model provides an opportunity to study the role of stroma in thyroid biology. Further, the cellular and molecular map generated here was exploited to predict potential communication between cell types (Figure 2E). The *in silico* interaction network, we hope, will help to develop hypotheses for testing the role of microenvironment in thyroid gland differentiation.

The current study provides the first single-cell atlas of mouse thyroid organoid, yet it has certain limitations. Firstly, the number of thyrocytes with mature characteristics (*Tg*+ *Tpo*+ *Slc5a5*+) profiled in the study are rather low, approximately 1.3 % of the total profiled cells. In future, it would be of interest to enrich for mature thyrocytes, potentially by exclusively profiling EGFP+ population in late stages of differentiation. Secondly, we lack an *in vivo* scRNA-Seq reference of endocrine differentiation into thyroid follicles. Currently, a single-cell atlas of adult thyroid gland for zebrafish (40) and humans (47), and bulk RNA-Seq data of mouse thyroid progenitors (48) is present in literature. However, we lack a detailed atlas of anterior foregut endoderm undergoing thyroid differentiation at cellular resolution. Without an *in vivo* reference, we are missing information on potential factors that are absent *in vitro* but could potentially improve the maturation of the organoid.

Further, in future, it would be of interest to establish a multi-dimensional atlas of the thyroid organoid and organ by simultaneous profiling of chromatin accessibility and protein levels along with transcriptome, potentially by utilizing protocols that forgo enzymatic dissociation (49) to reduce genetic perturbation associated with cell collection. Such integrative dataset would allow development of gene regulatory networks active during distinct stages of thyroid differentiation. This could elucidate redundancy and feedback loops present in the system, allowing better identification of gene targets for improving the robustness of differentiation protocol.

## 4 Material and Methods

### 4.1 ESC culture for maintenance and differentiation

A2Lox.Cre_TRE-Nkx2.1/Pax8_Tg-Venus mouse ESCs (Figure 1A) were cultured and were differentiated as described previously by Antonica, 2012 (9). Briefly, embryoid bodies, generated by hanging drops culturing of ESCs (1,000 cells per drop) were collected after 4 days and embedded in growth-factor-restricted Matrigel (MTG; BD Biosciences); 50µl MTG drops (containing around 20 EBs) were plated into 12-well plates. Embryoid bodies were differentiated using a differentiation medium initially supplemented with 1µg/ml of Doxycyclin (Sigma) for 3 days, followed by two weeks of maturation by using differentiation medium containing 0.3 nmol of 8-Br-cAMP (BioLog) (Figure 1B). Cells were collected and fixed (PFA 4%) at day 21 in order to confirm the thyroid differentiation (Figure 1C).

### 4.2 FACS sorting of mouse thyroid organoid

At differentiation days: 7, 12, 17 and 21, organoids cultured in MTG drops were washed twice with Hanks’s balanced salt solution (HBSS, containing calcium and magnesium; Invitrogen) and incubated for 30 min at 37 °C in a solution (1 ml per well) containing 10 U/ml of dispase II (Roche) and 125 U/ml of collagenase type IV (Sigma) in HBSS. Cells were dissociated and re-suspended manually by using a P1000 pipette and collected in a 15-ml Falcon tube (3 wells per tube, in duplicates, per time point). Then the enzymes were inactivated by adding 10% FBS and cells were centrifuged at 500g for 3 min. Cells were rinsed twice with HBSS following centrifugation settings described above. Finally, the organoids were incubated with TrypLE Select solution (Thermo Fisher, 12563011) for 10 min at 37°C, in order to dissociate then to single cells. Following dissociation, TrypLE solution was inactivated with 10% FBS, and single cells pelleted by centrifugation at 500 g for 3 min. The pellet was resuspended in 500 µl of HBSS, and the solution was filtered using a 30 µm cell filter (pluriSelect, 43-50030-03). In order to remove dead cells, calcein violet (Thermo Fisher, C34858) was added (final concentration of 1 µM) to the cell suspension and it was incubated at room temperature for 20 min. The single-cell preparation was sorted with the appropriate gates, including excitation with 405 nm laser for identification of alive cells (calcein+) and excitation with 488 nm for thyroid (Tg-EGFP+) cells (Figure 1D). FACS was performed through 100 µm nozzle.

### 4.3 Single-cell RNA-Sequencing of the mouse thyroid organoid

For single-cell RNA-seq of the thyroid in vitro organoids using the 10× Genomics platform, cell suspension was prepared as mentioned above from the thyroid organoids collected at differentiation days 7, 12, 17 and 21. For each stage, cells were dissociated and collected by FACS as described above. 4000 alive cells for each stage were collected in a single tube. For day 7, cells were collected irrespective of EGFP expression, while for days 12, 17 and 21, 50% EGFP+ and 50% EGFP-cells were collected. Sequencing library from the collected cells was prepared and sequenced following the previously described protocol (40).

In order to build the reference for Cell Ranger, mouse genome (GRCm38) and gene annotation (Ensembl 87) were downloaded from Ensembl and the annotation was filtered with the “mkgtf “ command of Cell Ranger (options: “-attribute = gene_biotype:protein_coding-attribute = gene_biotype:lincRNA -attribute = gene_biotype:antisense”). Within, the mouse genome and filtered gene annotation, the last exon of *Pax8* gene was masked. The last exon for *Pax8* was introduced and labeled as exogenous Nkx2.1_Pax8 sequence (10x Chromium 3’ RNA Sequencing kit only sequences ∼100 bp upstream of the polyA tail. For endogenous *Pax8*, this would correspond to sequences in the 3’ UTR. For exogenous *Pax8*, this would correspond to sequences in the last exon, as the exogenous *Pax8* does not harbor a 3’ UTR after the coding sequence. The exogenously provided sequence is a bicistronic construct containing *Nkx2*.*1* and *Pax8*. Hence, levels of exogenous *Pax8* reflect levels of the bicistronic *Nkx2*.*1, Pax8* construct.) The modified genome sequence and annotation were then used as input to the “mkref “ command of Cell Ranger to build the appropriate Cell Ranger Reference.

### 4.4 Cluster analysis of ScRNA-Seq data

The raw data generated from Cell Ranger pipeline was processed using Scanpy 1.6.0 package (25) by following the recommended pipeline. Briefly, filtering was performed on raw data to remove low quality cells by filtering cell profiles that had high mitochondrial gene content, low number of captured unique molecules or expressed low number of genes (Figure 1E). To remove duplicates, cell profiles with high UMI or genes were further excluded (Figure 1E). After quality control, raw data was log-normalized, cell cycle score was calculated. The effect of cell cycle, library size and mitochondrial counts was regressed out. Finally, the data was scaled for downstream analysis. Highly variable genes were utilized to compute principal component analysis and neighborhood graph (UMAP). For clustering, the first forty principal components were utilized as they covered 90% of the standard deviation as accessed by elbow plot. Further, a resolution of 0.4 was used for leiden clustering. Differential gene expression analysis was performed for each cluster using Wilcoxon rank-sum test to identify marker genes.

To subcluster the thyroid lineage, distributions of normalized gene expressions were plotted for *Tg, Slc5a5, Tpo, Hhex, Foxe1, Nkx2-1* and *Pax8*. For each gene, cells expressing the gene above 0.5 times normalized expression were labeled as expressing cells.

### 4.5 Gene Ontology (GO) Analysis

GO term analysis was performed using Enrichr (50) by submitting the complete rank gene list obtained using Wilcoxon rank-sum test. Genes that were statistically enriched (False Discovery Rate (FDR) < 0.05) with a minimum log fold enrichment of 1.5 within a cluster were chosen for analysis. For visualization, statistically significant (p-value < 0.05) terms were selected from KEGG 2019 Human, WikiPathways 2019 Human, GO Biological Process 2018 and GO Molecular Function 2018.

### 4.6 Cell-cell interaction analysis of ScRNA-Seq data

Receptor-ligand interaction analysis is performed utilizing CellPhoneDB package (26). Input data from Scanpy object was extracted as outlined in CellPhoneDB guidelines. Prior to analysis, mouse gene names were converted to human gene names retrieved from BioMart (51). The statistical method was run using the default parameters. Dot plot was plotted using a manually selected list of ligand-receptor pairs displaying statistically significant interaction. For a detailed description of the terms, please refer to CellPhoneDB documentation.

### 4.7 Diffusion pseudotime analysis of ScRNA-Seq data

For single cell trajectory inference, diffusion pseudotime was computed following the standard analysis pipeline (52) provided by Scanpy package. Since a single lineage was selected for the analysis, number of branchings was set to 1 and epithelial progenitors were selected as root cells. To visualize the marker genes trajectory on trendline plot, plot.wishbone_marker_trajectory function was used provided by Wishbone package (53).

### 4.8 Pharmacological inhibition of TGF-beta pathway

In order to inhibit TGF-beta signaling, thyroid organoids were cultured in differentiation medium containing cAMP and SB431542 inhibitor (10µM; Peprotech) from day 22 to day 29 of differentiation protocol and the medium was changed every 48 h. At day 29, thyroid organoids were collected for gene expression, immunofluorescence and iodide organification.

### 4.9 Quantitative PCR (qPCR) analysis

qPCR was performed on cDNA generated from thyroid organoids at differentiation day 29. For this, total RNA was extracted from thyroid organoids by lysis using RLT Lysis buffer (Qiagen) + 1% 2-mercaptoethanol. Extracted RNA was isolated using RNeasy micro kit (Qiagen) according to the manufacturer’s instructions. Reverse transcription was done using Superscript II kit (Invitrogen) to generate cDNA. qPCR was performed on cDNA in triplicates using Takyon (Eurogentec) and CFX Connect Real-Time System (Biorad). Results are presented as linearized values normalized to the housekeeping gene, B2microglobulin and the indicated reference value (2-DDCt). The gene expression profile was obtained from three independent samples. Primers used were as follows: B2microglobulin: Fw 5’-GCTTCAGTCGTCAGCATGG-3’,Rv5’-CAGTTCAGTATGTTCGGCTTCC-3’; Nkx2.1: Fw 5’-GGCGCCATGTCTTGTTCT-3’, Rv 5’-GGGCTCAAGCGCATCTCA-3’;Pax8:Fw5’-CAGCCTGCTGAGTTCTCCAT-3’, Rv 5’-CTGTCTCAGGCCAAGTC CTC-3’; Foxe1: Fw 5’-GGCGGCATCTACAAGTTCAT-3’,Rv5’-GGATCTTGAGGAAGCAGTCG-3’;THSr:Fw5’-GTCTGCCCAATATTTCCAGGATCTA-3’, Rv 5’-GCTCTGTCAAGGCATCAGGGT-3’; Slc5a5 (Nis): Fw 5’-AGCTGCCAACACTTCCAGAG-3’, Rv 5’-GATGAGAGCAC CACAAAGCA-3’;Tg:Fw5’-GTCCAATGCCAAAATGATGGTC-3’,Rv5’-GAGAGCATCGGTGCTGTTAAT-3’;Tpo:Fw5’-ACAGTCACAGTTCTCCACGGATG-3’,Rv5’-ATCTCTATTGTTGCACGCCCC-3’; Duox2: Fw 5’ -AACGGCACTCTCTGACATGG-3’, Rv 5’ -GGCCCCATTACCTTTTTGCC-3’.

### 4.10 Immunofluorescence

For protein immuno-detection experiments, cells were fixed in 4% paraformaldehyde (Sigma) for 1 h at room temperature (RT) and washed three times in PBS. Cells were blocked in a solution of PBS containing 3% bovine serum albumin (BSA; Sigma), 5% horse serum (Invitrogen) and 0.3% Triton X-100 (Sigma) for 30 min at RT. The primary and secondary antibodies were diluted in a solution of PBS containing 3% BSA, 1% horse serum and 0.1% Triton X-100. Primary antibodies against Tg (rabbit anti-TG, A0251 Dako, 1:2,000), Nis (rabbit anti-NIS, a gift from N. Carrasco, 1:1,000) and Tg-I (mouse anti-TG-I, a gift from C. Ris-Stalpers, 1:100) were incubated overnight at 4°C followed by incubation with secondary antibodies (donkey anti-mouse and anti-rabbit IgG conjugated with DyLight-Cy3 and DyLight-647; 1:500; Jackson Immunoresearch) and Hoechst 33342 (1:1,000; Invitrogen) for 2 h at RT. Coverslips were mounted with Glycergel (Dako). The samples were imaged on Zeiss LSM 780 confocal microscope using a 32x magnification.

### 4.11 Iodide organification assay

Thyroid organoids at differentiation day 29, treated with cAMP or cAMP+SB431542, were initially washed with HBSS and incubated with 1 ml of an organification medium containing 1,000,000 c.p.m. per ml ^125^I (PerkinElmer) and 100 nM sodium iodide (Sigma) in HBSS for 2 h at 37 °C. After incubation, 1 ml of 4 mM MMI, was added to the cells and washed with ice-cold PBS. Then, cells were detached using 0.1% trypsin (Invitrogen) and 1 mM EDTA (Invitrogen) in PBS for 15 min. For iodide uptake quantification, cells were collected in polyester tubes and radioactivity was measured with a c-counter. Subsequently, proteins were precipitated twice by addition of 1 mg of gamma-globulins (Sigma) and 2 ml 20% TCA followed by centrifugation at 2,000 r.p.m. for 10 min, at 4°C and the radioactivity of protein-bound [^125^I] (PBI) was measured. Iodide organification was calculated as an iodide uptake/PBI ratio and the values expressed as a percentage. As control for iodide uptake and protein-binding measurements cells were also treated with 30µM sodium perchlorate (Nis inhibitor; NaClO4, Sigma-Aldrich)) and 2 µM methimazole (TPO inhibitor; MMI, Sigma-Aldrich), respectively. The experiments were performed in triplicates for each condition.

### 4.12 Statistical analysis

Gene expression levels are shown in fold changes and compared using one-way ANOVA with a post-hoc Turkey’s comparison test. Uptake and protein-bound levels are expressed as mean ± SD and compared by unpaired Student’s t-test.

## 6 Author Contributions

S.P.S. and M.R. conceptualized the project. M.R. and B.F. generated and maintained thyroid organoids. S.P.S. performed the single-cell RNA-Sequencing. S.E.E. performed the bioinformatics analysis. M.R. performed pharmacological manipulations and with help of B.F. performed the subsequent analysis. M.R., S.E.E. and B.F. wrote the first draft and S.C., S.P.S. edited the manuscript. S.C. and S.P.S. acquired funding for the project. All authors read and approved the final manuscript.

## 8 Acknowledgments

We thank members of the Costagliola and Singh lab for comments on the manuscript. We thank J.- M. Vanderwinden from the Light Microscopy Facility and Christine Dubois from the FACS facility for technical assistance at ULB. S.C. would like to acknowledge the support from FNRS as a Directeur de recherches.

## 9 Data Availability Statement

All NGS data (raw files and counts) are uploaded to GEO with accession number GSE163818 and reviewer token wlohckywbvixlqr. Scripts to analyze the data and raw images will be made available upon request.

## 11 Supplementary Material

**Supplementary table 1:** Differentially expressed marker genes. Marker genes display minimum 1.5 log-fold change and FDR (false discovery rate) of less than 0.05.

**Supplementary table 2:** Interacting ligand-receptor pairs from CellPhoneDB. In the significant means tab, insignificant pairs display zero mean expression, while the significant pairs are indicated with mean expression values.

## References

1. Santos-Ferreira TF, Borsch O, Ader M. Rebuilding the Missing Part-A Review on Photoreceptor Transplantation. Front Syst Neurosci (2017) 10: doi:10.3389/fnsys.2016.00105

2. Hayek A, King CC. Brief review: cell replacement therapies to treat type 1 diabetes mellitus. Clin Diabetes Endocrinol (2016) 2:4. doi:10.1186/s40842-016-0023-y

3. Fan Y, Winanto Ng S-Y. Replacing what’s lost: a new era of stem cell therapy for Parkinson’s disease. Transl Neurodegener (2020) 9:2. doi:10.1186/s40035-019-0180-x

4. Mattapally S, Pawlik KM, Fast VG, Zumaquero E, Lund FE, Randall TD, Townes TM, Zhang J. Human Leukocyte Antigen Class I and II Knockout Human Induced Pluripotent Stem Cell-Derived Cells: Universal Donor for Cell Therapy. J Am Heart Assoc (2018) 7: doi:10.1161/JAHA.118.010239

5. Riolobos L, Hirata RK, Turtle CJ, Wang P-R, Gornalusse GG, Zavajlevski M, Riddell SR, Russell DW. HLA Engineering of Human Pluripotent Stem Cells. Mol Ther (2013) 21:1232–1241. doi:10.1038/mt.2013.59

6. Sagar Grün D. Deciphering Cell Fate Decision by Integrated Single-Cell Sequencing Analysis. Annu Rev Biomed Data Sci (2020) 3:1–22. doi:10.1146/annurev-biodatasci-111419-091750

7. Kumar P, Tan Y, Cahan P. Understanding development and stem cells using single cell-based analyses of gene expression. Development (2017) 144:17–32. doi:10.1242/dev.133058

8. Griffiths JA, Scialdone A, Marioni JC. Using single-cell genomics to understand developmental processes and cell fate decisions. Mol Syst Biol (2018) 14: doi:10.15252/msb.20178046

9. Antonica F, Kasprzyk DF, Opitz R, Iacovino M, Liao X-H, Dumitrescu AM, Refetoff S, Peremans K, Manto M, Kyba M, et al. Generation of functional thyroid from embryonic stem cells. Nature (2012) 491:66–71. doi:10.1038/nature11525

10. Kurmann AA, Serra M, Hawkins F, Rankin SA, Mori M, Astapova I, Ullas S, Lin S, Bilodeau M, Rossant J, et al. Regeneration of thyroid function by transplantation of differentiated pluripotent stem cells. Cell Stem Cell (2015) 17:527. doi:10.1016/j.stem.2015.09.004

11. Nilsson M, Fagman H. Development of the thyroid gland. Development (2017) 144:2123–2140. doi:10.1242/dev.145615

12. Rousset B, Dupuy C, Miot F, Dumont J. Chapter 2 Thyroid Hormone Synthesis And Secretion. (2000). Available at: http://www.ncbi.nlm.nih.gov/pubmed/25905405

13. Fernández LP, López-Márquez A, Santisteban P. Thyroid transcription factors in development, differentiation and disease. Nat Rev Endocrinol (2015) 11:29–42. doi:10.1038/nrendo.2014.186

14. De Felice M, Di Lauro R. Thyroid Development and Its Disorders: Genetics and Molecular Mechanisms. Endocr Rev (2004) 25:722–746. doi:10.1210/er.2003-0028

15. Slack JMW. From Egg to Embryo. Cambridge University Press (1991). doi:10.1017/CBO9780511525322

16. Plachov D, Chowdhury K, Walther C, Simon D, Guenet JL, Gruss P. Pax8, a murine paired box gene expressed in the developing excretory system and thyroid gland. Development (1990) 110:643–51. Available at: http://www.ncbi.nlm.nih.gov/pubmed/1723950

17. Lazzaro D, Price M, de Felice M, Di Lauro R. The transcription factor TTF-1 is expressed at the onset of thyroid and lung morphogenesis and in restricted regions of the foetal brain. Development (1991) 113:1093–104. Available at: http://www.ncbi.nlm.nih.gov/pubmed/1811929

18. Zannini M, Avantaggiato V, Biffali E, Arnone MI, Sato K, Pischetola M, Taylor BA, Phillips SJ, Simeone A, Di Lauro R. TTF-2, a new forkhead protein, shows a temporal expression in the developing thyroid which is consistent with a role in controlling the onset of differentiation. EMBO J (1997) 16:3185–97. doi:10.1093/emboj/16.11.3185

19. Fagman H, Andersson L, Nilsson M. The developing mouse thyroid: Embryonic vessel contacts and parenchymal growth pattern during specification, budding, migration, and lobulation. Dev Dyn (2006) 235:444–455. doi:10.1002/dvdy.20653

20. Opitz R, Maquet E, Huisken J, Antonica F, Trubiroha A, Pottier G, Janssens V, Costagliola S. Transgenic zebrafish illuminate the dynamics of thyroid morphogenesis and its relationship to cardiovascular development. Dev Biol (2012) 372:203–16. doi:10.1016/j.ydbio.2012.09.011

21. Postiglione MP, Parlato R, Rodriguez-Mallon A, Rosica A, Mithbaokar P, Maresca M, Marians RC, Davies TF, Zannini MS, De Felice M, et al. Role of the thyroid-stimulating hormone receptor signaling in development and differentiation of the thyroid gland. Proc Natl Acad Sci U S A (2002) 99:15462–7. doi:10.1073/pnas.242328999

22. Milenkovic M, De Deken X, Jin L, De Felice M, Di Lauro R, Dumont JE, Corvilain B, Miot F. Duox expression and related H2O2 measurement in mouse thyroid: onset in embryonic development and regulation by TSH in adult. J Endocrinol (2007) 192:615–26. doi:10.1677/JOE-06-0003

23. Islam S, Zeisel A, Joost S, La Manno G, Zajac P, Kasper M, Lönnerberg P, Linnarsson S. Quantitative single-cell RNA-seq with unique molecular identifiers. Nat Methods (2014) 11:163–6. doi:10.1038/nmeth.2772

24. McInnes L, Healy J, Melville J. UMAP: Uniform Manifold Approximation and Projection for Dimension Reduction. (2018) Available at: http://arxiv.org/abs/1802.03426

25. Wolf FA, Angerer P, Theis FJ. SCANPY: large-scale single-cell gene expression data analysis. Genome Biol (2018) 19:15. doi:10.1186/s13059-017-1382-0

26. Efremova M, Vento-Tormo M, Teichmann SA, Vento-Tormo R. CellPhoneDB: inferring cell-cell communication from combined expression of multi-subunit ligand-receptor complexes. Nat Protoc (2020) 15:1484–1506. doi:10.1038/s41596-020-0292-x

27. Haerlingen B, Opitz R, Vandernoot I, Trubiroha A, Gillotay P, Giusti N, Costagliola S. Small Molecule Screening in Zebrafish Embryos Identifies Signaling Pathways Regulating Early Thyroid Development. Thyroid (2019)551861. doi:10.1089/thy.2019.0122

28. Trapnell C. Defining cell types and states with single-cell genomics. Genome Res (2015) 25:1491–8. doi:10.1101/gr.190595.115

29. Oguchi H, Kimura S. Multiple Transcripts Encoded by the Thyroid-Specific Enhancer-Binding Protein (T/EBP)/Thyroid-Specific Transcription Factor-1 (TTF-1) Gene: Evidence of Autoregulation*. Endocrinology (1998) 139:1999–2006. doi:10.1210/endo.139.4.5933

30. di Gennaro A, Spadaro O, Baratta MG, De Felice M, Di Lauro R. Functional Analysis of the Murine Pax8 Promoter Reveals Autoregulation and the Presence of a Novel Thyroid-Specific DNA-Binding Activity. Thyroid (2013) 23:488–496. doi:10.1089/thy.2012.0357

31. De Felice M, Di Lauro R. Thyroid development and its disorders: genetics and molecular mechanisms. Endocr Rev (2004) 25:722–46. doi:10.1210/er.2003-0028

32. Mio C, Grani G, Durante C, Damante G. Molecular defects in thyroid dysgenesis. Clin Genet (2020) 97:222–231. doi:10.1111/cge.13627

33. Castanet M, Marinovic D, Polak M, Léger J. Epidemiology of thyroid dysgenesis: the familial component. Horm Res Paediatr (2010) 73:231–7. doi:10.1159/000284386

34. Maenhaut C, Christophe D, Vassart G, Dumont J, Roger P., Opitz R. Ontogeny, Anatomy, Metabolism and Physiology of the Thyroid. (2000). Available at: http://www.ncbi.nlm.nih.gov/pubmed/25905409

35. Mincione G, Di Marcantonio MC, Tarantelli C, D’Inzeo S, Nicolussi A, Nardi F, Donini CF, Coppa A. EGF and TGF-B1 Effects on Thyroid Function. J Thyroid Res (2011) 2011:431718. doi:10.4061/2011/431718

36. López-Márquez A, Fernández-Méndez C, Recacha P, Santisteban P. Regulation of Foxe1 by Thyrotropin and Transforming Growth Factor Beta Depends on the Interplay Between Thyroid-Specific, CREB and SMAD Transcription Factors. Thyroid (2019) 29:714–725. doi:10.1089/thy.2018.0136

37. Cortijo C, Gouzi M, Tissir F, Grapin-Botton A. Planar Cell Polarity Controls Pancreatic Beta Cell Differentiation and Glucose Homeostasis. Cell Rep (2012) 2:1593–1606. doi:10.1016/j.celrep.2012.10.016

38. Flasse L, Yennek S, Cortijo C, Barandiaran IS, Kraus MR-C, Grapin-Botton A. Apical Restriction of the Planar Cell Polarity Component VANGL in Pancreatic Ducts Is Required to Maintain Epithelial Integrity. Cell Rep (2020) 31:107677. doi:10.1016/j.celrep.2020.107677

39. Bader E, Migliorini A, Gegg M, Moruzzi N, Gerdes J, Roscioni SS, Bakhti M, Brandl E, Irmler M, Beckers J, et al. Identification of proliferative and mature B-cells in the islets of Langerhans. Nature (2016) 535:430–4. doi:10.1038/nature18624

40. Gillotay P, Shankar M, Haerlingen B, Sema Elif E, Pozo-Morales M, Garteizgogeascoa I, Reinhardt S, Kränkel A, Bläsche J, Petzold A, et al. Single-cell transcriptome analysis reveals thyrocyte diversity in the zebrafish thyroid gland. EMBO Rep (2020) n/a:e50612. doi:10.15252/embr.202050612

41. Lupulescu A. “Ultrastructural pathology of the thyroid gland,” in Ultrastructure of Endocrine Cells and Tissues, ed. P. M. Motta (Boston, MA: Springer US), 286–301. doi:10.1007/978-1-4613-3861-1_25

42. Min S, Kim S, Cho S-W. Gastrointestinal tract modeling using organoids engineered with cellular and microbiota niches. Exp Mol Med (2020) 52:227–237. doi:10.1038/s12276-020-0386-0

43. Karpus ON, Westendorp BF, Vermeulen JLM, Meisner S, Koster J, Muncan V, Wildenberg ME, van den Brink GR. Colonic CD90+ Crypt Fibroblasts Secrete Semaphorins to Support Epithelial Growth. Cell Rep (2019) 26:3698-3708.e5. doi:10.1016/j.celrep.2019.02.101

44. Lahar N, Lei NY, Wang J, Jabaji Z, Tung SC, Joshi V, Lewis M, Stelzner M, Martín MG, Dunn JCY. Intestinal Subepithelial Myofibroblasts Support in vitro and in vivo Growth of Human Small Intestinal Epithelium. PLoS One (2011) 6:e26898. doi:10.1371/journal.pone.0026898

45. Brügger MD, Valenta T, Fazilaty H, Hausmann G, Basler K. Distinct populations of crypt-associated fibroblasts act as signaling hubs to control colon homeostasis. PLOS Biol (2020) 18:e3001032. doi:10.1371/journal.pbio.3001032

46. Jolly LA, Novitskiy S, Owens P, Massoll N, Cheng N, Fang W, Moses HL, Franco AT. Fibroblast-Mediated Collagen Remodeling Within the Tumor Microenvironment Facilitates Progression of Thyroid Cancers Driven by Braf V600E and Pten Loss. Cancer Res (2016) 76:1804–1813. doi:10.1158/0008-5472.CAN-15-2351

47. Han X, Zhou Z, Fei L, Sun H, Wang R, Chen Y, Chen H, Wang J, Tang H, Ge W, et al. Construction of a human cell landscape at single-cell level. Nature (2020) 581:303–309. doi:10.1038/s41586-020-2157-4

48. Ikonomou L, Herriges MJ, Lewandowski SL, Marsland R, Villacorta-Martin C, Caballero IS, Frank DB, Sanghrajka RM, Dame K, Kandula MM, et al. The in vivo genetic program of murine primordial lung epithelial progenitors. Nat Commun (2020) 11:635. doi:10.1038/s41467-020-14348-3

49. Eski SE, Dubois C, Singh SP. Nuclei Isolation from Whole Tissue using a Detergent and Enzyme-Free Method. J Vis Exp (2020) doi:10.3791/61471

50. Kuleshov M V., Jones MR, Rouillard AD, Fernandez NF, Duan Q, Wang Z, Koplev S, Jenkins SL, Jagodnik KM, Lachmann A, et al. Enrichr: a comprehensive gene set enrichment analysis web server 2016 update. Nucleic Acids Res (2016) 44:W90–W97. doi:10.1093/nar/gkw377

51. Smedley D, Haider S, Ballester B, Holland R, London D, Thorisson G, Kasprzyk A. BioMart - biological queries made easy. BMC Genomics (2009) 10:22. doi:10.1186/1471-2164-10-22

52. Haghverdi L, Büttner M, Wolf FA, Buettner F, Theis FJ. Diffusion pseudotime robustly reconstructs lineage branching. Nat Methods (2016) 13:845–8. doi:10.1038/nmeth.3971

53. Setty M, Tadmor MD, Reich-Zeliger S, Angel O, Salame TM, Kathail P, Choi K, Bendall S, Friedman N, Pe’er D. Wishbone identifies bifurcating developmental trajectories from single-cell data. Nat Biotechnol (2016) 34:637–45. doi:10.1038/nbt.3569

